# Natural variation in root exudation of GABA and DIMBOA impacts the maize root endosphere and rhizosphere microbiomes

**DOI:** 10.1101/2021.11.05.467511

**Authors:** Peng Wang, Lucas Dantas Lopes, Martha G. Lopez-Guerrero, Karin van Dijk, Sophie Alvarez, Jean-Jack Riethoven, Daniel P. Schachtman

## Abstract

Root exudates are important for shaping root-associated microbiomes. However, studies on a wider range of metabolites in exudates are required for a comprehensive understanding about their influence on microbial communities. We identified maize inbred lines that differ in exudate concentrations of DIMBOA and GABA using a semi-hydroponic system. These lines were grown in the field to determine the changes in microbial diversity and gene expression due to varying concentrations of DIMBOA and GABA in exudates using 16S rRNA amplicon sequencing and metatranscriptomics. Results showed individual and interaction effects of DIMBOA and GABA on the rhizosphere and root endosphere β-diversity, most strongly at the V10 growth stage. The main bacterial families affected by both compounds were *Ktedonobacteraceae* and *Xanthomonadaceae*. Higher concentrations of DIMBOA in exudates affected the rhizosphere metatranscriptome, enriching for KEGG pathways associated with plant disease. This study validated the use of natural variation within plant species as a powerful approach for understanding the role of root exudates on microbiome selection. We also showed that a semi-hydroponic system can be used to identify maize genotypes that differ in GABA and DIMBOA exudate concentrations under field conditions. The impact of GABA exudation on root-associated microbiomes was shown for the first time.

## Introduction

Root exudation is an important strategy used by plants to interact with soil microbial communities [1]. The rhizosphere is the area around plant roots influenced by root secretion [2]. Root exudates are the main compounds secreted in the rhizosphere and includes a diversity of carbon-containing primary and secondary metabolites such as amino acids, sugars, organic acids and hormones [3], which serve as energy and carbon sources for the heterotrophic soil microbiota [4, 5]. In turn, soil microbial activity release nutrients for plant growth through organic matter decomposition, nitrogen (N) fixation, metal chelation, phytohormones production and also support plant health by alleviating abiotic stresses and suppressing pathogens [6–9]. Many factors have been shown to drive the amount and composition of exudates secreted into the rhizosphere [10, 11], including plant species, developmental stages, abiotic and biotic stresses, but little is known about the intra-species variation in root exudation or the precise role of each specific exudate compound [12].

Plant secondary metabolites including hormones are important in the selection of the root-associated microbiomes [13]. To determine the effect of exudates on microbiome composition studies have mainly been conducted using mutants of model plants, in which the production of a particular root exudate is abolished [13–15]. For example, a tomato mutant impaired in the exudation of the phytohormone strigolactone led to significantly decreased arbuscular mycorrhizal fungi colonization, which may negatively impact water and nutrient acquisition by the plant host [16, 17]. In two studies using *Arabidopsis* mutants [18, 19], the secondary metabolite coumarin was shown to be involved in shaping the microbiome under iron deficiency and inhibiting two fungal pathogens [18, 19]. Flavonoids, which are plant-derived rhizosphere signaling molecules, have very well-known roles in signaling between legume roots and rhizobia [20]. The compound 2,4-dihydroxy-7-methoxy-2H-1,4-benzoxazin-3(4H)-one (DIMBOA) is a benzoxazinoid found in the *Poaceae* family including maize and wheat. It plays roles in defense against herbivores [21]. DIMBOA has also been characterized for its effects on the root and rhizosphere microbiomes [14, 22, 23]. In maize mutants, benzoxazinoids (BXs) affected fungal and bacterial community structure in the root and rhizosphere of maize in a time-dependent manner [23].

Gamma-aminobutyric acid (GABA) is another secondary metabolite found in high concentrations in many plant species [24–26]. GABA is a non-proteinogenic amino acid found in animals [27], and also in some bacterial species [28]. GABA controls quorum sensing in *Agrobacterium* [29]. GABA is also involved in plant growth, development and stress responses, is present in root exudates [30] and may be transported to the rhizosphere by ALMT1 [31]. Despite the many roles that GABA plays in planta, the interaction between root exudates containing GABA and the root-associated microbiomes is unexplored.

At the present time there are limited numbers of root exudate mutants available in maize and other crop plants that would allow for the characterization of the influence of exudates in the composition and diversity of soil microbial communities. Therefore, in this study we used the natural variation in root exudate composition identified across different maize inbred lines to determine the impact of DIMBOA and GABA on the root endosphere and rhizosphere microbiomes. To our knowledge this is the first study to use a natural variation approach for studying the influence of root exudates on root-associated microbiomes. The natural variation found in plant genotypes has long been an important source of new diversity for agricultural crops [32]. Previous reports also suggested that natural variation may be present in maize exudates based on leaf concentrations of benzoxazinoids [32] and other metabolites [33].

Nine maize lines were selected from a large maize diversity panel [34, 35] based on different concentrations of DIMBOA and GABA in the root exudates of maize seedlings growing in an aseptic glass bead semi-hydroponic system. DIMBOA was chosen to validate this natural variation approach because it has been shown to affect the microbiome [14, 23, 36]. GABA was chosen because its role has been recognized as important in multiple processes in roots and as a signaling molecule [27]. Bacterial community composition was profiled using 16S rRNA amplicon sequencing and metatranscriptomics to understand how DIMBOA impacts the functional characteristics in the prokaryotic and fungal communities. The central questions investigated were: 1) Can changes in the microbiome be detected using natural variation in DIMBOA and GABA root exudates? 2) Is there an interaction between DIMBOA and GABA in shaping the bacterial communities? 3) How do the functional characteristics of microbial communities change due to DIMBOA in root exudates? Our study confirms the importance of root exudates in shaping the root-associated microbes and validates the use of plant natural variation in studying these processes. Natural variation will be important for elucidating the complexity of responses in the rhizosphere microbiome due to root exudates.

## Methods Summary

More details can be found in the Supplemental Methods. We selected nine maize genotypes belonging to the Goodman & Buckler association panel[34, 35], based on their concentrations of GABA and DIMBOA in the root exudates. We used a semi-hydroponic system to collect exudates in 1 mM CaCl_2_ for 2 hours and analyzed the concentrations of GABA and DIMBOA in the different genotypes through LC-MS. The nine maize lines selected were planted in a field experiment (40.86° N, 96.61° W) randomized in blocks, and samples from rhizosphere and root endosphere were collected at the V5, V10 and R2 growth stages. The sampling method for the root endosphere and rhizosphere was previously described [37]. In addition to measuring DIMBOA and GABA concentration using the semi-hydroponics system, we also used the supernatant of rhizosphere samples collected from the field to measure the concentration of the two compounds through LC-MS. The rhizosphere and root endosphere DNA were extracted using the MoBio PowerSoil-htp 96 well kit and the MagMax Plant DNA isolation kit, respectively. PCR amplifications of the 16S rRNA gene were performed using the 515F and 806R primer pair [38] with a dual-index method [39, 40]. The libraries were sequenced with 300 bp paired-end kit on an Illumina MiSeq. A subset of six genotypes including Ames10248, Ames27140, Ames27171, B73, PI587128, and NSL65873 were selected for the metatranscriptomics analysis at the V10 stage. The rhizosphere RNA was extracted with the Qiagen RNeasy PowerSoil Total RNA Kit (Cat No. 12866-25), and sequencing was performed with the 75 bp paired-end kit on the Illumina NextSeq500 platform. Data processing for the 16S rRNA gene reads was performed as described previously [37, 41] using QIIME (v1.9.1) [42], USEARCH and UPARSE softwares [43, 44], to generate a 97% identity OTU table taxonomically classified using the RDP database [45]. The mRNA-seq data was processed to trim the paired-end reads, filter sequences by length and quality using Trimmomatic v0.38 [46], remove maize contaminating reads aligning against the B73 genome sequence using the bowtie v2.3 software [47], removing rRNA sequences using the Infernal v1.1.2 software [48], assembly the transcripts using Trinity v2.8 [49], and generating a transcript count table using bowtie v2.3 and R software [50, 51]. After transcripts classification in different microbial groups using Kaiju 1.7 software [52], only archaeal, bacterial and fungal transcripts were kept. Finally, transcripts were annotated to the KEGG prokaryotic database using USEARCH [44] and to the COG/Pfam/CDD protein databases with RPSBlast (version 2.4.0) [53]. Alpha and beta-diversity analyses were done based on the rarefied OTU tables. The Bray-Curtis index was used for the unconstrained and constrained PCoA using the *capscale() function* and PerMANOVA using the *adonis()* function in ‘vegan’ package [54]. The concentrations of DIMBOA and GABA were used as quantitative variables in the multivariate analyses. Welch’s t-test was used to identify bacterial families with significantly different relative abundance between DIMBOA and GABA levels using the STAMP software [55]. For the metatranscriptomics analysis, the prokaryotic and fungal transcript table were analyzed independently through PCoA and PerMANOVA using the Bray-Curtis matrices. The prokaryotic and fungal taxa significantly changing in gene expression between DIMBOA levels were identified using the LEfSe software and edgeR package [56, 57]. Differences due to GABA concentration were not analyzed because only one genotype of each GABA level was sequenced (metatranscriptomics). KEGG enrichment analysis was performed in R software [58, 59]. In addition, Mantel tests were performed between the Bray-Curtis distance matrices of the 16S rRNA gene OTU table and the transcripts count table, as well as between the taxonomy tables (at family level) classified based on 16S rRNA amplicon or metatranscriptomics data.

## Results

### Changes in microbial diversity and community structure associated with different concentrations of GABA and DIMBOA in root exudates

Nine maize lines were chosen for these analyses (Fig. 1). The DIMBOA concentrations in exudates of three lines were classified as low and five lines were classified as high (Fig. 1A). The average GABA concentrations in exudates of two lines were classified as low and five lines as high (Fig. 1B). The exudate concentrations of the genotypes classified as low were significantly different to those classified as high for both DIMBOA and GABA.

**Figure 1.**
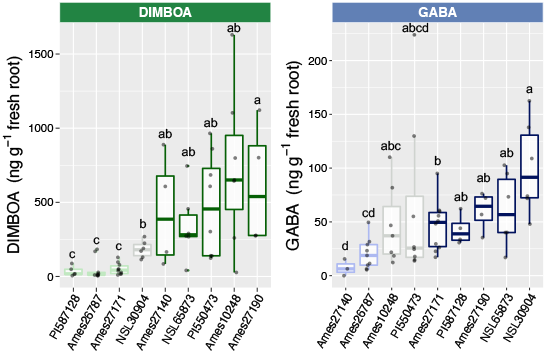
DIMBOA and GABA concentrations in the root exudates of nine genotypes collected from a semi-hydroponic system. (A) Concentrations of DIMBOA normalized by the root fresh weight in the nine genotypes. (B) Concentrations of GABA normalized by root fresh weight. Within the box plot, the line and in the boxes represent median, top and bottom of boxes represent first and third quartiles, respectively, and whiskers indicate 1.5 interquartile range. The letters on the top of the error bars indicate results of statistical analysis using the Kruskal-Wallis test. Different letters indicate significant differences (*p* < 0.05). The x axis indicates the accession number of the genotype. Two levels of DIMBOA and GABA (low and high) indicated by lighter green or blue as low level and darker green or blue for higher concentration level were classified based on significant differences between the genotypes.

Principal coordinate analysis (PCoA) and PerMANOVA showed that the compartment was the most important factor affecting bacterial community structure in this study (*p* < 0.001; R^2^ = 0.2), followed by growth stages (*p* < 0.001; R^2^ = 0.1) (Fig. 2A). Moreover, the interaction between compartment and growth stage had a significant (*p* < 0.001; R^2^ = 0.05) impact on bacterial community structure. Constrained PCoA was used to remove the effect of blocks in order to reduce the influence of spatial variation. There were significant differences between the two plant compartments (root endosphere *vs*. rhizosphere) (*p* < 0.001) and the three growth stages in each compartment (V5, V10 and R2) (Supplemental Fig. 1). The bacterial α-diversity was significantly higher in the rhizosphere of maize lines with low DIMBOA concentrations, but was not affected by the different GABA concentrations in root exudates (Supplemental Fig. 1).

**Figure 2.**
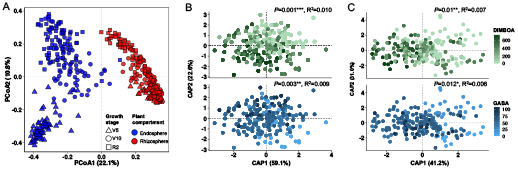
Bacterial β-diversity/community structure of the nine genotypes used in the field experiment according to growth stage, plant compartment, and the semi-hydroponic concentration of DIMBOA and GABA in the root exudates. (A) Principal coordinate analysis (PCoA) using the Bray-Curtis distance matrix shows the separation of samples based on growth stage and plant compartment. (B) CAP analysis showing the effect of DIMBOA and GABA concentration in root exudates on the bacterial community structure of rhizosphere samples. (C) CAP analysis showing the effect of DIMBOA and GABA concentration in root exudates on the bacterial community structure of root endosphere samples. Numerical values of DIMBOA and GABA concentrations were used for the ordination and to perform the PerMANOVA factoring out the effect of block and time in the model simultaneously. The PerMANOVA *p*-value and R^2^ are shown above each graph.

Further analyses were completed on the rhizosphere and endosphere separately because the bacterial community composition of the two compartments was significantly different. A constrained PCoA and PerMANOVA showed that both DIMBOA and GABA concentrations in exudates had a significant influence on the structure of rhizosphere and root endosphere bacterial communities (Fig. 2B-C). The natural variation in these two different exudates among maize inbred lines revealed that there was an interaction (*p* = 0.006) between these two exudates in shaping the rhizosphere bacterial community composition. To further explore the nature of the interaction, we split the 16S rRNA amplicon data for the maize lines into high and low exudate concentrations of DIMBOA and GABA and tested for significance under the different scenarios. The results showed that changes in the rhizosphere bacterial β-diversity associated with different GABA concentrations were only significant when DIMBOA levels in exudates were high (Supplemental Fig. 2A). In contrast, the rhizosphere bacterial β-diversity associated with the different DIMBOA exudate concentrations was significant at both GABA exudate concentrations (Supplemental Fig. 2B). In the root endosphere, the bacterial β-diversity associated with different GABA exudate concentrations were significant at both DIMBOA concentrations (Supplemental Fig. 2C), while the β-diversity associated with different DIMBOA concentrations were significant only when concentrations of GABA were low (Supplemental Fig. 2D). Shifts in α-diversity in the rhizosphere were only observed due to changes in GABA concentrations in the high DIMBOA lines (Supplemental Fig. 2E-F). Changes in α-diversity in the endosphere due to exudate concentrations were observed in the low levels of DIMBOA and GABA (Supplemental Fig. 2G-H). The interaction between exudates highlights the complexity of their influence on the bacterial communities.

To determine how developmental changes in maize may play a role in the temporal changes in the rhizosphere and endosphere, the data were separated into V5, V10 and R2 growth stages in each compartment. DIMBOA exerted significant effects on rhizosphere bacterial β-diversity at all stages of development (Supplemental Fig. 3A), but only significantly impacted the root endosphere at R2 (*p* = 0.007) (Supplemental Fig. 3C). The exudate concentrations of GABA significantly influenced microbial community composition at V10 (*p* = 0.003) for the rhizosphere and the root endosphere (*p* = 0.05) (Supplemental Fig. 3B and D). The interaction between DIMBOA and GABA was significant at V10 (*p* = 0.006; R^2^ = 0.035) and R2 (*p* = 0.03; R^2^ = 0.028) in the rhizosphere and only significant in the endosphere at V10 (*p* = 0.03; R^2^ = 0.024). At the different stages where we measured significant impacts of each exudate compounds on the β-diversity, the variation in the community composition explained by each factor (GABA and DIMBOA) was between 2.2% and 3.9% (Supplemental Fig. 3).

The maize lines used in this study were chosen based on root exudate concentrations in a semi-hydroponic system. However, the concentrations of these compounds were also measured in rhizosphere samples from field grown plants at V10 and R2. The concentrations of DIMBOA and GABA in the rhizosphere supernatant was also used to assess their influence on bacterial community structure. The different DIMBOA concentrations in the rhizosphere supernatant significantly affected the rhizosphere β-diversity at V10 and R2, while GABA concentrations was only significant at V10 (Supplemental Fig. 4A). The percentage of variation in the microbiome due to DIMBOA and GABA in the rhizosphere was 5% and 5.6%, respectively. DIMBOA concentrations in the rhizosphere did not significantly impact the root endosphere microbiome at any stage, while GABA concentrations affected the endosphere microbiome at the R2. The percentage of variation accounted for GABA and DIMBOA in the root endosphere was ≤ 3.6%. These analyses using exudate concentrations measured in rhizosphere samples indicated that the semi-hydroponics data provided a good estimation of how the different inbred lines would exude these compounds in the field.

### Changes in specific microbial taxa between the different root exudates concentrations

We further investigated changes in relative abundance of bacterial taxa between maize lines exuding different levels of DIMBOA and GABA. When analyzing all the growth stages together, 35 bacterial families were significantly different in relative abundance between maize genotypes exuding high or low levels of DIMBOA in the rhizosphere (Fig. 3A). Among them, *Xanthomonadaceae* and *Ktedonobacteraceae* showed the largest differences between the two levels and were enriched in the genotypes exuding more DIMBOA. In the root endosphere only eight bacterial families were significantly different in relative abundance between DIMBOA concentrations (Fig. 3B). *Oxalobacteraceae* showed the largest difference in relative abundance, followed by *Xanthomonadaceae* – both were enriched in genotypes exuding more DIMBOA. In contrast to DIMBOA, more bacterial families were significantly different in relative abundance in the root endosphere (32) than in the rhizosphere (14) between the lines that differed in GABA concentrations in root exudates (Fig. 3C-D). Similar to the results with DIMBOA, *Xanthomonadaceae* and *Ktedonobacteraceae* were also enriched in the genotypes exuding more GABA in the rhizosphere (Fig. 3C). The bacterial family with the largest difference in relative abundance between GABA concentrations in the root endosphere was *Alcaligenaceae*, enriched in the genotypes exuding a low concentration of this compound, followed by *Streptomycetaceae* and *Ktedonobacteraceae*, which were in higher relative abundance in the maize genotypes exuding more GABA (Fig. 3D). *Ktedonobacteraceae* and *Xanthomonadaceae* were the bacterial families responding most to the high levels of GABA and DIMBOA in the root exudates in both rhizosphere and endosphere.

**Figure 3.**
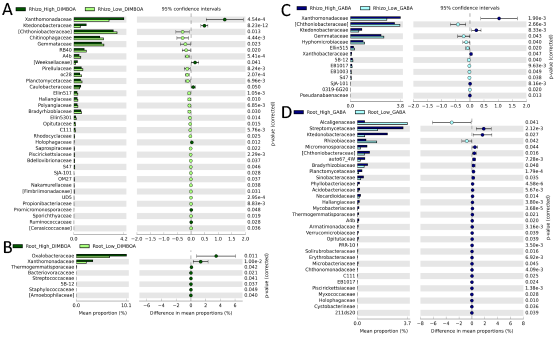
Differences in the relative abundance of bacterial families between maize genotypes classified as high and low DIMBOA or GABA exuders. (A) Bacterial families significantly affected by DIMBOA levels (high vs. low) in the rhizosphere. (B) Bacterial families significantly affected by DIMBOA levels (high vs. low) in the root endosphere. (C) Bacterial families significantly affected by GABA levels (high vs. low) in the rhizosphere. (D) Bacterial families significantly affected by GABA levels (high vs. low) in the root endosphere. The Welch’s t-test was used to identify the bacterial families showing significant differences between DIMBOA and GABA concentrations in exudates using the software Statistical Analysis of Metagenomic Profiles (STAMP). Only the families with corrected *p-*value ≤ 0.05 (Bonferroni) were shown in the bar graphs. The classification of genotypes as low and high exuders of DIMBOA and GABA was according to Fig.1.

The analysis at three times during development revealed growth stage specific differences in relative abundance of bacterial families in maize lines with high and low DIMBOA exudate concentrations. The number of bacterial families showing significant differences in relative abundance between genotypes exuding different DIMBOA concentrations in the rhizosphere *vs*. root endosphere were 16 *vs*. 3 (V5), 23 *vs*. 5 (V10), and 34 *vs*. 7 (R2), respectively (Supplemental Fig. 5A). The numbers of bacterial families with significantly different relative abundance between genotypes exuding different GABA levels in the root endosphere *vs*. rhizosphere was 4 *vs*. 6 (V5), 16 *vs*. 26 (V10), and 8 *vs*. 32 (R2), respectively (Supplemental Fig. 5B). The families *Ktedonobacteraceae* and *Xanthomonadaceae* were enriched in the lines with high GABA and high DIMBOA exudate concentrations in almost all growth stages and compartments. The separation of stages highlighted several other families that were enriched at one or two growth stages. At the V5 stage *Pseudomonadaceae, Spingobacteriaceae, Weeksellaceae* and *Enterobacteriaceae* were enriched in the rhizosphere and *Enterobacteriaceae* was enriched in the endosphere of maize lines with high DIMBOA in exudates. In the lines with high GABA concentrations *Planctomycetaceae* and *Isopharaceae* were also enriched at V5, *Anaeroplasmataceae* at V10 and R2, and *Sphingomonadaceae* at R2.

### Changes in rhizosphere metatranscriptome associated with different exudate concentrations

PCoA and PerMANOVA indicated that the metatranscriptome of both prokaryotic and fungal communities were significantly different between genotypes with different exudate concentrations of DIMBOA (*p* = 0.003 for prokaryote and *p* = 0.01 for fungi). These results show that the taxonomic structure and gene expression of the microbial communities were influenced by the DIMBOA concentrations in root exudates. A total of 463 prokaryotic and 1,043 fungal genes were differentially expressed due to high vs. low DIMBOA concentrations (*FDR corrected p* < 0.05), (Supplemental Table 1 and 2).

The differences in gene expression were further explored by assessing the KEGG pathways that were significantly affected by DIMBOA concentrations. For prokaryotic genes, seven KEGG pathways were enriched due to higher concentrations of DIMBOA including bacterial secretion systems, biofilm formation, plant-pathogen interaction and MAPK signaling pathways; while three KEGG pathways were enriched due to lower DIMBOA levels including the biosynthesis of antibiotics, microbial metabolism and metabolic pathways (Fig. 5A). For fungi, three KEGG pathways associated with sugar metabolism were enriched in rhizosphere samples due to higher levels of DIMBOA, *i*.*e*. galactose metabolism, starch and sucrose metabolism, and amino and nucleotide sugar metabolism (Fig. 5B).

We also tested for correlations between the transcript count table and the OTU table (16S rRNA amplicon) on the same rhizosphere samples at the V10 stage. A significant positive correlation between the two datasets (*p* = 0.027) was found (Mantel test), although the Spearman’s correlation coefficient was relatively low (ρ = 0.29). When comparing the taxonomy at the family level based on the 16S rRNA amplicon data and metatranscriptomics data, no significant correlation was detected. These results indicate that differences in microbial community structure are consistent with differences in gene expression, but the taxa (families) changing in abundance and in gene expression were not the same.

Independent of the maize inbred exudate concentrations, the most dramatic difference in taxonomic classification at the phylum level between the 16S rRNA gene and metatranscriptomics data was for Actinobacteria and Firmicutes, which were over-represented in the metatranscriptomics data, indicating the high expression of genes from these phyla in the rhizosphere compared to other phyla (Fig. 6A). In contrast, Planctomycetes, Chloroflexi and Verrucomicrobia were under-represented in metatranscriptome data as compared to the 16S rRNA data for DIMBOA. Basidiomycota transcripts dominated the fungal metatranscriptome for DIMBOA, followed by Ascomycota (Fig. 6B).

Eight bacterial families, including *Chitinophagaceae, Planctomycetaceae, Ktedonobacteraceae, Bacillaceae, Polyangiaceae, Gemmatimonadaceae, Flavobacteriaceae*, and *Chlamydiaceae*, showed significant changes in gene expression between low and high DIMBOA levels (Supplemental Fig. 6A), but among those families only three - *Chitinophagaceae, Planctomycetaceae* and *Ktedonobacteraceae* – also showed changes in relative abundance based on the 16S rRNA gene data at this same stage (V10) (Supplemental Fig. 5A). The changes in both relative abundance and gene expression of these three bacterial families indicate they were the most impacted by DIMBOA exudates. The gene expression of 17 fungal genera were influenced by the different DIMBOA levels (Supplemental Fig. 6B). Among the fungal genera with significant changes in gene expression due to DIMBOA, fourteen were enriched when exudate concentrations of DIMBOA were high including *Saitoella, Fusarium, Mortierella, Bipolaris, Melampsora, Aspergillus, Oidiodendron, Phialocephala, Gelatoporia, Talaromyces, Hypoxylon, Glomus, Grifola* and *Exophiala*, while only three genera were enriched when DIMBOA concentrations were low, *i*.*e. Fibulorhizoctonia, Rhizoctonia* and *Botryobasidium* (Supplemental Fig. 6B).

## Discussion

In this study we analyzed the effects of variation in the root exudation of DIMBOA and GABA on the belowground microbial communities associated with maize roots. The natural variation in GABA and DIMBOA exudates were first analyzed in a growth chamber semi-hydroponic system (Fig. 1) and later the differences in exudate concentrations were confirmed by analyzing the supernatant of the solutions we used to release rhizosphere soil from roots (Supplemental Fig. 4). Analyzing the natural variation among genotypes of a plant species is an approach that may have certain advantages compared to using plant mutants of root exudation processes [14, 23, 36]. For example, generating mutants may be time consuming, particularly in crop plants, because it may be difficult to identify the genes to mutate to alter production and exudation pathways. It may also be possible to identify a larger number of genotypes with variation in the concentration of a root exudate. Multiple genotypes also broaden the applicability of the study, increase the statistical power and enhance confidence in the results because multiple genetic backgrounds are being used. This is particularly important when the study aims to infer the global effects at the plant species level. On the other hand this approach may introduce more noise due to unrelated genotypes which could also reduce statistical power to detect significant effects. In the case of maize, there is a large amount of genotypic variation within the species and analysis of a mutation in a single inbred line may be subject to background genotype effects as demonstrated for DIMBOA mutants [36, 60].

While other exudates not studied in this work may also impact the composition of the bacterial microbiome the approach used here was based on Permutational Multivariate Analysis of Variance (PerMANOVA). This analysis provides a measure of the percent contribution of each compound studied to the total variation. In this study GABA and DIMBOA contributed 2 – 5% of the total variation while other factors such as spatial heterogeneity which was removed from the analysis and growth stage generally have a larger impact on total variation. Typically statistical models do not include all factors because many are unknown. Eventhough additional cofactors could be added to our model to improve the fit, our results clearly showed the two exudates characterized have significant effects on the bacterial communities and highlight that the contribution of the exudates is small but significant. Our findings of the significant impacts of GABA and DIMBOA on bacterial communities strengthens the conclusion that the use of natural variation is a viable and productive method to study the impacts of specific root exudates on the root associated bacterial communities.

DIMBOA was selected to validate this approach because previous studies using a mutational approach indicated that this compound affects the root-associated microbial communities of wheat and maize [14, 23, 36, 61]. It has also been shown to have important functions for plants such as chelation of soil iron and defense against plant herbivores [62, 63]. In addition to the changes in microbial community structure analyzed by previous studies, we assessed the metatranscriptome changes of the rhizosphere microbiome due to different concentrations of this exudate which has not been previously reported.

Two studies assessing the changes in the maize root-associated microbiomes using mutants in *bx* genes detected a higher percentage of variation (determined by PerMANOVA R^2^) explained by DIMBOA levels than we observed [14, 23]. A third study found no significant changes in the microbial β-diversity [36]. Our study analyzed multiple lines exuding different concentrations of DIMBOA and we found smaller but significant changes in β-diversity, which may reflect differences that would be expected from crops in the field. The three studies mentioned above found few or no changes in microbial α-diversity due to DIMBOA root content [23, 36] or exudation [14]. In contrast, we found higher species diversity under lower concentrations of DIMBOA at all three growth stages (Supplementary Fig. 1). Two previous studies showed that bacterial community structure was more affected by DIMBOA than the fungal communities which were also affected [14, 23]. Our metatranscriptomics data are in concordance with previous findings at the DNA level in that we found the prokaryotic transcriptional profile is more affected by DIMBOA than were fungal transcripts (Fig. 4).

**Figure 4.**
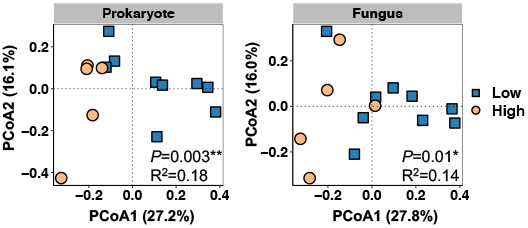
Changes in the rhizosphere metatranscriptome of prokaryotic and fungal communities based on the different DIMBOA concentrations in the root exudates. Principal coordinate analysis (PCoA) based on Bray-Curtis distance matrices of transcripts counts were performed using 17 rhizosphere samples collected from six genotypes. PerMANOVA was used to test the effects of DIMBOA levels in root exudates on the prokaryotic and fungal metatranscriptomes. The *p-*value and R^2^ are shown for each graph. Only a subset of genotypes showing significant differences in the exudate concentration of DIMBOA were selected for the metatranscriptomics analysis. The maize lines used for this analysis included: PI587128, Ames26787 and Ames27171 that were classified as low DIMBOA exuders, B73 and Ames10248 that were classified as high DIMBOA exuders.

The significant changes in the prokaryotic metatranscriptome point toward an increase in the expression of genes associated with pathogenesis in the lines exuding more DIMBOA. In addition to the obvious “plant-pathogen interaction”, the KEGG pathways “MAPK signaling pathway”, and “bacterial secretion system” enriched under high DIMBOA levels are all potentially associated with pathogenic plant-microbe interactions (Fig. 5A). The MAPK cascades are essential for plant innate immune responses and many microbial pathogens are known to perturb these cascades in order to evade the host immune system [64, 65]. The bacterial secretion systems may be associated with microbe-microbe interactions such as microbial competition or horizontal gene transfer (conjugation, transformation), but most types (I-VII) of secretion systems are used by phytopathogenic bacteria to produce adhesion and motility structures, and to secrete toxins, cell-degrading enzymes, and effectors to infect and evade the immune system to cause plant diseases [66]. Abiotic stress conditions also repress the *Arabidopsis* immune system and enrich for bacteria with the immune-activating versions of the flagellin epitope flg22, typical of phytopathogens [67]. Since DIMBOA is usually associated with signaling response to a biotic stress (insect herbivory), the plant immune system may be impacted thereby increasing the abundance and activity (gene expression) of phytopathogenic bacteria. The relative abundance analysis supported this inference by showing a significantly higher abundance of *Xanthomonadaceae* in root endosphere and rhizosphere of lines with higher DIMBOA exudation levels (Fig. 3A). *Xanthomonas* spp. are widely known as pathogens of crops such as maize [68].

**Figure 5.**
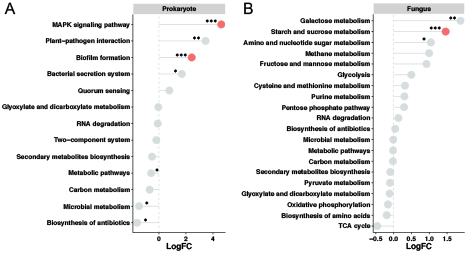
Functional pathways affected by DIMBOA concentrations in root exudates based on the metatranscriptome data for rhizosphere prokaryotic and fungal communities. (A) KEGG pathway enrichment analysis showing the pathways significantly enriched or depleted under high DIMBOA levels in root exudates based on the prokaryotic transcripts. (B) KEGG pathway enrichment analysis showing the pathways significantly enriched or depleted under high DIMBOA levels in root exudates based on the fungal transcripts. The values on the x axis indicate the log2–fold transformation of the relative percentages of each KEGG Onthology (KO) enriched under high/low DIMBOA relative to the total percentage of each KO within the entire dataset. The pathways with at least three significantly affected transcripts in (A) bacteria and five in (B) fungus are shown. But the pathways that were significantly different compared to the background using Fisher test are indicated using symbols “***” *p*<0.001, “**” *p*<0.01 and “*” *p*<0.05. Some of the grey circles are highlighted with red color, which indicates significant differences after the Benjamini-Hochberg (BH) *p*-value correction. The maize lines used for this analysis included: PI587128, Ames26787 and Ames27171 that were classified as low DIMBOA exuders, B73 and Ames10248 that were classified as high DIMBOA exuders.

**Figure 6.**
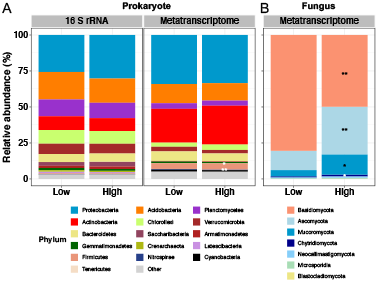
Bar charts showing the relative abundance of prokaryotic and fungal phyla according to the 16S rRNA and/or metatranscriptomics approaches. (A) Comparison between the proportion of bacterial phyla as determined by 16S rRNA gene sequencing and the proportion of transcripts assigned to bacterial phyla as determined by metatranscriptomics. The asterisks indicate significant differences in the proportion of specific phyla between the high and low DIMBOA levels. (B) Comparison between the proportion of transcripts assigned to fungal phyla in the low versus high DIMBOA levels. Asterisks indicate significant differences in specific phyla between the low and high levels of DIMBOA. The symbols “**” and “*” indicate *p*<0.01 and *p*<0.05 by Wilcoxon rank sum test, respectively. The maize lines used for metatranscriptomics analysis included: PI587128, Ames26787 and Ames27171 that were classified as low DIMBOA exuders, B73 and Ames10248 that were classified as high DIMBOA exuders; For the relative abundance of phyla based on the 16S rRNA gene sequencing, all the genotypes classified as low and high DIMBOA according to Fig. 1 were included.

The enrichment of the KEGG pathway “biofilm formation” in the lines exuding more DIMBOA is probably associated with more bacterial colonization [69]. In contrast, the lines exuding less DIMBOA were enriched in the “biosynthesis of antibiotics”, a mechanism used for biocontrol of plant pathogens by beneficial plant-growth promoting rhizobacteria (PGPR) [9]. The other bacterial family strongly affected by DIMBOA exudate concentrations was *Ktedonobacteraceae*, which was enriched in maize lines exuding high DIMBOA at all growth stages in the rhizosphere compartment, and at V10 in the root endosphere (Supplemental Fig. 5A). The gene transcripts associated with this family also increased under high DIMBOA (Supplemental Fig. 6A). Bacteria from this family are understudied [70], but a genome survey suggested that they are able to produce a wide range of secondary metabolites, including antibiotics [71]. Therefore, the higher relative abundance of this family under higher DIMBOA exudate concentration could possibly be in response to higher pathogen presence/activity. Another bacterial family enriched under high DIMBOA in the root endosphere was *Oxalobacteraceae*, which was previously shown to improve maize performance under N deprivation [72]. Since in our experiment the plants were not N-limited, it is possible that the increase in *Oxalobacteraceae* abundance is associated with plant stress. On the fungal side, we observed an increase in sugar metabolism pathways associated with the lines exuding more DIMBOA (Fig. 5B). Sugars (mainly sucrose) are important molecules used in molecular plant-fungal interactions [73], and by biotrophic fungal pathogens [74]. Together, these results suggest that genotypes producing lower DIMBOA concentration in the exudates are less susceptible to soil pathogens than lines with higher DIMBOA production. The lower rhizosphere α-diversity found during the whole growing season in the lines exuding higher DIMBOA levels could be associated with stressful conditions that favor pathogens establishment in the rhizosphere.

GABA is known to be an important compound for plant developmental and physiological processes, it accumulates in plant tissues in response to biotic and abiotic stresses [75], and has been proposed as an endogenous signaling molecule [76]. GABA also negatively regulates anion efflux via ALMT transporters [25, 27] which have also been shown to mediate cellular efflux and influx of GABA [31]. In contrast to DIMBOA, the effects of GABA exudates on the plant-associated microbial communities are unknown except for one study that suggested a correlation between GABA added to soil and some microbial groups [77]. Here we showed that the concentration of GABA in root exudates also affects the rhizosphere and root endosphere microbiomes of maize.

In contrast to DIMBOA that affected the rhizosphere β-diversity at all growth stages and the root endosphere at the R2 stage, GABA affected both rhizosphere and root endosphere β-diversity only at the V10 stage (Supplemental Fig. 3). Changes in the root-associated microbiomes along plant developmental stages were observed in our present study with maize, as well as in previous studies with *Arabidopsis, Avena barbata*, wheat and sorghum [6, 61, 78, 79]. It is likely that changes in exudate composition shapes the microbial community dynamics during plant development, where shifts in the concentration of sugars, hormones, and secondary metabolites are important in this selective process [6, 78, 79]. GABA may also play a role in developmental change in the microbiome in that the greatest effect of GABA on the microbiomes was at the V10 stage.

The largest changes in relative abundance of bacteria in maize lines differing in GABA exudation was also observed at V10, whereas the largest differences due to DIMBOA exudation were observed at the R2 stage (Supplemental Fig. 5B). However, the main bacterial families changing in relative abundance between maize lines due to GABA were *Xanthomonadaceae* and *Ktedonobacteraceae*, which were enriched on the lines exuding higher levels of GABA. The increase in *Xanthomonadaceae* under high GABA concentrations could be associated with a repressed immune system due to the connection between biotic stress and GABA accumulation [75], while the enrichment of *Ktedonobacteraceae* could be a plant-driven response to control the increase of plant pathogens [71]. However, elevated levels of GABA in *Arabidopsis* reduce the virulence of *Pseudomonas syringae* [80]. Interestingly, the root colonization of the biocontrol strain *Pseudomonas protegens* CHAO was shown to increase through GABA production, which is initiated by the GacA/GacS two component system [81]. Thus, one hypothesis is that GABA participates in plant-microbe signaling and activates the root colonization of biocontrol bacteria, which is a question that needs further investigation. It is noteworthy that the Actinobacterial family *Streptomycetaceae* widely known for producing a vast array of antibiotics [82] was also enriched on the root endosphere of genotypes exuding more GABA at the R2 stage.

An alternative reason for why those two bacterial families were enriched on lines exuding higher concentrations of both DIMBOA and GABA is the interactive effect these exudates may have on the bacterial communities. Significant interaction effects of DIMBOA and GABA on the rhizosphere β-diversity were found at the V10 and R2 stage, and at the V10 stage in the root endosphere (Supplemental Fig. 3). Moreover, the effect of DIMBOA on the rhizosphere and endosphere β-diversity was larger in the lines exuding less GABA, while the effect of GABA on the rhizosphere β-diversity was larger in the lines exuding more DIMBOA, and lower in the endosphere β-diversity of lines exuding more DIMBOA (Supplemental Fig. 2). Although the biological interactions between the two compounds on the root-associated bacterial communities studied here needs further confirmation, our results suggest that different exudate compounds may act synergistically on the selection of the root-associated bacterial communities. The interactive effects of exudates highlight the complexity of the roles that exudates play in shaping microbiomes. Therefore, for a comprehensive and mechanistic understanding of how the plant microbiome is shaped it will be important to consider the interactive effects of the multiple root exudates, and how different combinations and concentrations can affect the associated microbiomes. This study shows that natural variation is an appropriate and potentially useful approach to studying the impact of root exudates on microbial communities. The complexity and interactive effects of exudates were highlighted and we demonstrated that the exudation of GABA along with DIMBOA play roles in shaping the rhizosphere and endosphere microbiomes.

## Acknowledgements

We thank Stephanie Futrell, Bertrand Devilbiss for technical assistance and Dr. Ling Xu for advice on statistical analysis of the metatranscriptome data. The work was supported by the National Science Foundation EPSCOR (award OIA-1557417) to the Center for Root and Rhizobiome Innovation.

## Author contribution

PW and DPS planned and designed the research. PW, MGL_G and DPS performed the lab experiments. PW and DPS executed the field work and analyzed the data. LDL wrote the manuscript draft and conducted some analyses. PW and DPS edited the manuscript. SA did the exudate analysis. JJR helped with the transcript analysis. KVD supervised the identification of maize lines.

## Data availability

The data can be found at NCBI under the accession numbers PRJNA728503 (16S rRNA) and PRJNA728686 (mRNA-seq).

## Supplemental Materials and Methods

### Plant materials

The nine maize inbred lines used in this study were Ames 10248 (GA209), Ames 27140 (NC260), Ames 27190 (T234), PI550473 (B73), Ames 26787 (H49), Ames 27171 (NC350), NSL 30904 (4226), PI 587128 (H84) and NSL 65873 (L317). They were identified by screening 240 maize lines that belong to the Goodman & Buckler association panel [34, 35].

### Semi-hydroponic system for exudate assessment

The nine maize lines were selected based on the amount of GABA and DIMBOA in their root exudates. To collect the root exudates, plants were grown for two weeks in an aseptic glass bead semi-hydroponic system and were intermittently irrigated with sterile 0.25 x Hoagland’s nutrient solution to ensure roots remained moist and were not hypoxic. Root exudates were collected for 2 hour in 1 mM CaCl_2_. Solutions were then frozen, freeze dried (Labconco FreeZone Dryer), resuspended in 2% formic acid, desalted using a Solid Phase Extraction cartridges (Oasis MCX Cartridge, Waters Company) and analyzed by targeted LC-MS using standards. The number of replicates for each genotype with exudates collected were Ames 10248 (n = 7), Ames 27140 (n=4), Ames 27190 (n=4), PI550473 (n=8), Ames 26787 (n=9), Ames 27171 (n = 10), NSL 30904 (n=6), PI 587128 (n=5) and NSL 65873 (n=6).

### Field experiments

The nine maize genotypes were planted on May 9, 2018, at a University of Nebraska - Lincoln experimental field (40.86° N, 96.61° W). The experiment was a randomized block design with eight blocks, with 0.91m between rows. Plants were sampled on May 31^th^ at V5 growth stage, July 2^nd^ at V10 growth stage and July 25^th^ at R2 growth stage. No fertilizer, herbicides or pesticides were applied, and no additional irrigation was needed. The previous crop was soybean. Rainfall was 27.7 cm from May 9 to July 25, 2018. The average bulk soil moisture on the three sampling collection dates were 20.1% at V5, 19.3% at V10, and 13.1% at R2 stages. The collection method for the root endosphere and rhizosphere, and the 16S rRNA gene sequencing has been previously described [37]. Amplicon sequencing of the V4 region of the 16S rRNA gene was completed on 72 rhizosphere samples and 68 endosphere samples at V5 stage, 72 rhizosphere and 71 endosphere samples at V10 stage, and 69 rhizosphere and 69 endosphere samples at R2 stage. For the nucleic acid extraction from rhizosphere soil, roots were removed from soil, and the soil adhering to the roots was collected and immediately placed on dry ice and stored at -80°C until extraction.

### Quantification of DIMBOA and GABA collected from rhizosphere field samples

Rhizosphere samples from the field collected in 50 mL tubes containing phosphate buffer [37] were placed on ice, brought to the lab and centrifuged to pellet the rhizosphere soil. The supernatant was carefully transferred into a new 50 mL tube and the rhizosphere soil was weighed. The supernatant was frozen at -80°C, freeze dried at -50°C (Labconco FreeZone Dryer), desalted, and analyzed as described above. A total of 47 samples from the V10 stage and 64 samples from the R2 stage were collected. The weight of the rhizosphere soil pellet was used to normalize the metabolite data.

### Nucleic acid extraction and 16S rRNA gene and metatranscriptomics sequencing

The rhizosphere DNA was extracted using the MoBio PowerSoil-htp 96 well kit, while the root endosphere DNA was extracted using the Applied Biosystems (ThermoFisher Scientific) MagMax Plant DNA isolation kit. The PCR amplification of the 16S rRNA gene libraries were performed using the 515F and 806R primer pair [38] with a dual-index method [39,40]. The libraries were sequenced with 300 bp paired-end kit on an Illumina MiSeq. The Qiagen RNeasy PowerSoil Total RNA Kit (Cat No. 12866-25) was used to extract RNA from rhizosphere. Library preparation was performed using the NuGen Ovation Prokaryotic-seq protocol with insert size of 200 bp, and 150 ng input per sample. The 75 bp paired-end next-generation sequencing was done with Illumina NextSeq500 platform. A subset of six genotypes including Ames10248, Ames27140, Ames27171, B73, PI587128, and NSL65873 were selected for the metatranscriptomics analysis. Three replicates of rhizosphere samples were collected from each selected genotype at V10 stage. Among these 18 samples, one replicate from genotype B73 was deemed to be an outlier using hierarchical clustering and PCA analysis and removed for subsequent analysis.

### Bioinformatics analyses of sequencing reads

Data processing for the 16S rRNA gene reads was performed as described previously [37, 41] using QIIME (v1.9.1) [42], USEARCH and UPARSE software [43, 44]. An OTU table at the 97% identity was constructed using the UPARSE pipeline. Taxonomic assignments were done with the ribosomal database project classifier (RDP) [45]. Mitochondrion and plastid sequences, as well as bacterial and archaeal unclassified sequences were removed. OTUs with less than 4 counts in all the samples and those only present in less than three samples [12] were removed. Root endosphere and rhizosphere samples were rarefied to 7,051 reads per sample. When endosphere and rhizosphere were analyzed separately, they were rarefied to 7,000 and 12,799 reads, respectively.

Data processing of the mRNA-seq data included eight steps: (1) paired-end reads were trimmed using Trimmomatic v0.38 [46] with a 2 bp sliding window cutting bases off the start and end of the reads below a quality threshold of Q20. Reads with length less than 35bp were removed. (2) Reads that originated from maize were removed after aligning against the B73 genome sequence using the bowtie v2.3 software [47]. (3) The rRNA sequences were removed using the Infernal v1.1.2 software [48]. (4) Transcripts were assembled using Trinity v2.8 with a minimal contig length of 100 bp [49]. (5) A transcript count table was generated for differential abundance analysis by aligning all samples against the assembled transcripts using bowtie v2.3. The raw expression counts were extracted using the alignment file (gff format) and the counts were merged into a combined data matrix in R software using the “GenomicFeatures”, “GenomicAlignments” and “Rsamtools” packages [50, 51]. (6) Transcripts were then classified into bacteria, archaea, virus, fungi or other microbial eukaryotes using Kaiju 1.7 software [52] with the nr_euk database (June 2019) and only archaeal, bacterial and fungi transcripts were used. (8) Transcripts were annotated by aligning their protein translations to the KEGG prokaryotic database using USEARCH [44] and to the COG/Pfam/CDD protein databases with RPSBlast (version 2.4.0) [53]. The first hit of each transcript obtained in KEGG Orthology (KO) IDs were used to lookup Gene Ontology (GO) IDs, pathways, and enzyme commission (EC) numbers. In the case that no KO ID was found via KEGG alignments, the hits against COG/Pfam/CDD were used.

### Statistical analyses

Bray-Curtis distance matrix based on the OTU table was used for the unconstrained principal coordinate analysis (PCoAs) and canonical analysis of principal coordinates (CAP) using the *capscale() function* in ‘vegan’ package [54]. Permutational multivariate analysis of variance (PerMANOVA) were performed in R software using the *adonis()* function from the ‘vegan’ package to assess the significance of each experimental factor and their interaction using a maximum of 999 permutations [83]. The concentrations of DIMBOA and GABA in exudates measured in the semi-hydroponic system was normalized by root weight and the averages of each genotype were used as a quantitative variable in the multivariate analyses. The concentrations of DIMBOA and GABA measured from field samples were normalized by the rhizosphere pellet weight and the value of each sample was used as a quantitative variable in the multivariate analyses. The ordination of the PCoA and CAP were generated using the two axis explaining the largest amount of data variability with the ggplot2 package [84]. Differential relative abundance of bacterial families between DIMBOA and GABA levels in root exudates were performed using the Welch’s t-test in STAMP software [55]. Differences were considered significant if *p* < 0.05 after the Bonferroni *p*-value correction. For that, the maize genotypes were previously classified in low or high levels of GABA and DIMBOA exudation according to the semi-hydroponics data. The Kruskal-Wallis test was used to identify the maize genotypes significantly different (*p* < 0.05) in DIMBOA and GABA concentrations in the root exudates, and based on that they were classified in the high or low levels of each compound.

For the metatranscriptomic data analysis, two Bray-Curtis dissimilarity matrices were generated for the prokaryotic and fungal genes from their normalized count tables, and used for PCoA and PerMANOVA. In addition, all the transcripts were classified at the microbial phylum and family levels using kaiju v1.7 [52] to assess which taxa are affected in gene expression by the concentrations of DIMBOA. The analysis of changes in transcriptomics profile according to differences in GABA concentration was not performed due to the low number of replicates (only 1 genotype classified as high vs. 1 genotype classified as low). The prokaryotic and fungal phyla and families significantly changing in gene expression between high and low DIMBOA levels were identified using the LEfSe software [56]. Differential abundance analysis was also performed on the annotated transcripts using the edgeR package [57]. KEGG enrichment analysis through Fisher test for number of KO IDs was performed in R software (version 3.6.3) [58, 59]. A Mantel test was performed between the Bray-Curtis distance matrices of the 16S rRNA gene OTU table and the transcripts count table to determine whether bacterial/archaeal community structure and the metatranscriptome were correlated, using the PAST software [85]. In addition, a Mantel test was also performed between taxonomy tables (at family level) classified based on 16S rRNA amplicon or metatranscriptomics data.

