## Supplementary material for "Natural variation in root exudation of GABA and DIMBOA impacts the maize root endosphere and rhizosphere microbiomes": Supplmental Figures

### Supplemental Figures

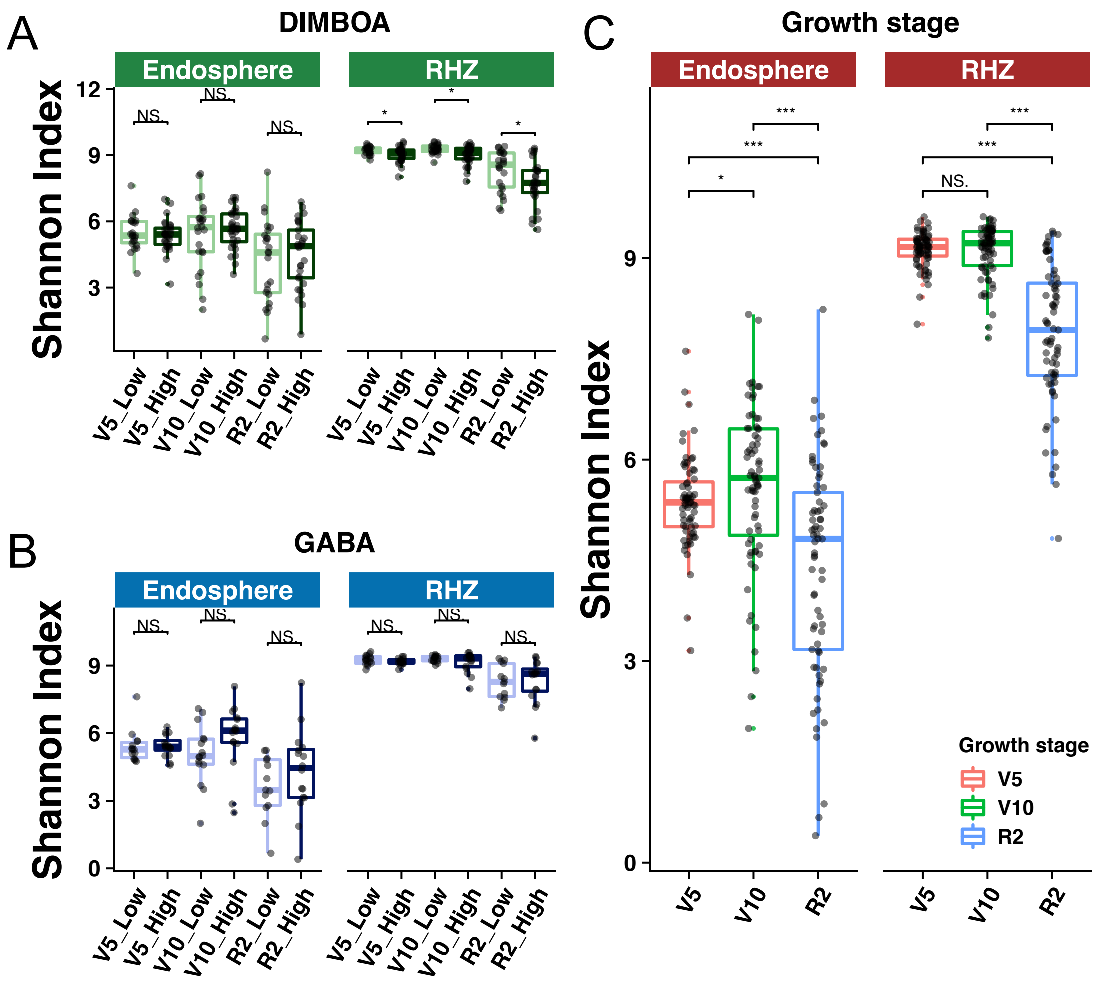


**Supplemental Figure S1. Shifts in α-diversity between maize genotypes with different DIMBOA and GABA levels in root exudates and between plant developmental stages.** (A) Differences in species diversity calculated by Shannon index in the root endosphere and rhizosphere bacterial communities due to DIMBOA levels in root exudates at different plant growth stages. (B) Differences in species diversity calculated by Shannon index in the root endosphere and rhizosphere bacterial communities due to GABA levels in root exudates along different plant growth stages. (C) Changes in species diversity (Shannon index) between the three plant developmental stages analyzed in the root endosphere and rhizosphere bacterial communities. Lines in the boxes represent the medians, top and bottom of boxes represent first and third quartiles, whiskers indicate 1.5 interquartile range, and dots outside of the whiskers range indicate the outliers. The symbols “*”, “**” and “***” indicate significant differences at $0.01<p<0.05$, $p<0.01$ and $p<0.001$ by Wilcoxon rank sum test, respectively. RHZ = rhizosphere samples; Endosphere = root endosphere samples.

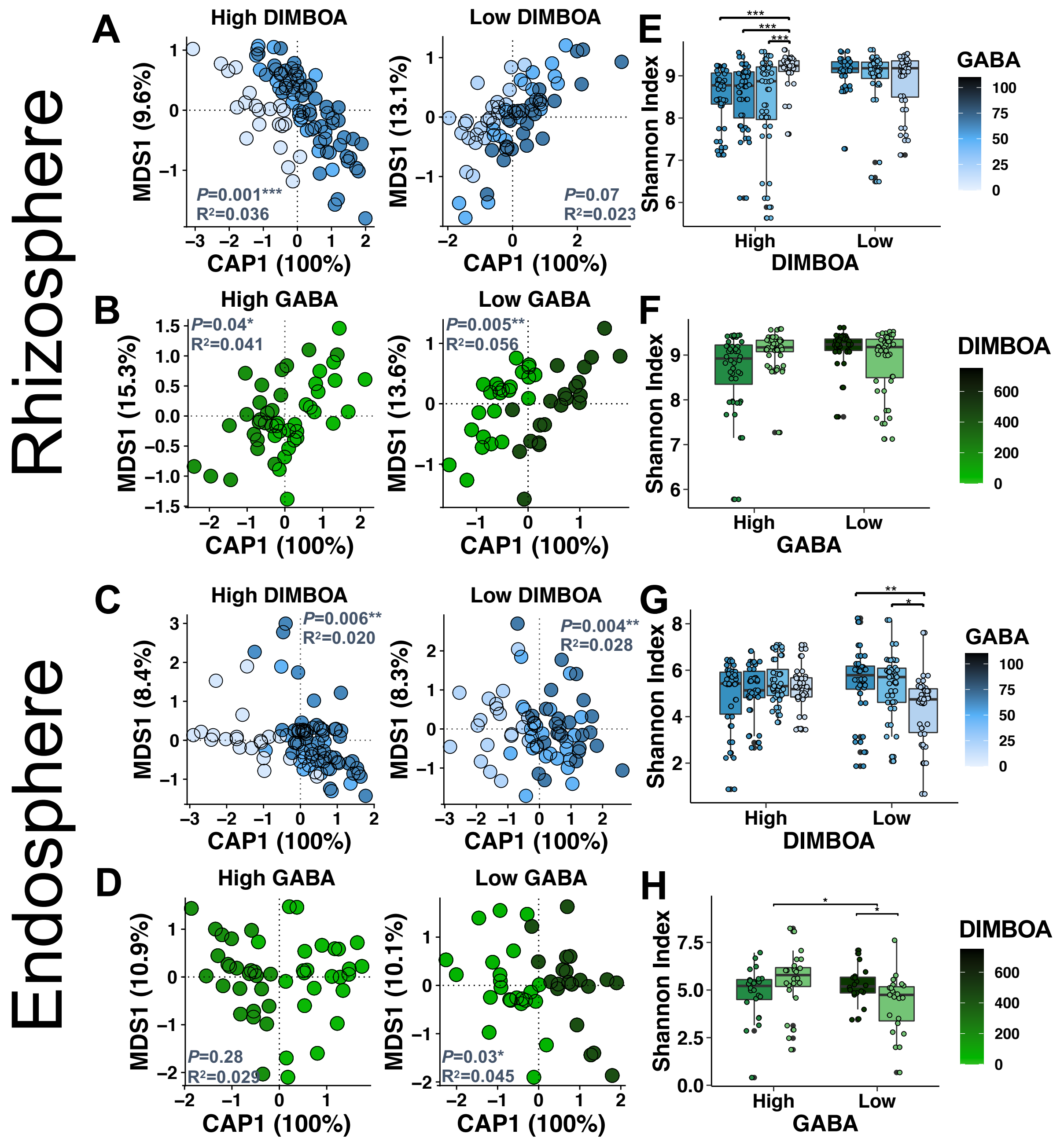


**Supplemental Figure S2. Effect of DIMBOA and GABA levels in root exudates on the bacterial community shaping within each level of the two compounds.** (A-D) Effect of DIMBOA on the β-diversity of bacterial communities in maize lines with high or low exudate concentrations of GABA and *vice-versa*. (A) CAP analysis showing the effect of GABA on the rhizosphere bacterial community structure in the high and low DIMBOA concentrations. (B) CAP showing the effect of DIMBOA on the rhizosphere bacterial community structure in the high or low GABA concentrations. (C) CAP showing the effect of GABA on the root endosphere bacterial community structure in the high and low DIMBOA concentrations. (D) CAP analysis showing the effect of DIMBOA on the root endosphere bacterial community structure in the high or low GABA concentrations. (E-H) Effect of DIMBOA on the α-diversity of bacterial communities in the high or low level of GABA and *vice-versa*. (E) Rhizosphere bacterial species diversity (Shannon index) due to GABA in maize lines with high or low exudate concentrations of DIMBOA levels. (F) Rhizosphere bacterial species diversity (Shannon index) due to DIMBOA in maize lines with high or low exudate concentrations of GABA levels. (G) Root endosphere bacterial species diversity (Shannon index) due to GABA in maize lines with high or low exudate concentrations of DIMBOA. (H) Root endosphere bacterial species diversity (Shannon index) due to DIMBOA in maize lines with high or low exudate concentrations of GABA. The symbols "*", "**" and "***" indicate significant differences at $p<0.05$, $p<0.01$ and $p<0.001$ by Wilcoxon rank sum test, respectively. The $p$-value and $R^2$ in each CAP plot indicate the results of PerMANOVA constrained for DIMBOA or GABA and factoring out the effect of "Block" from the field experiment in the model.

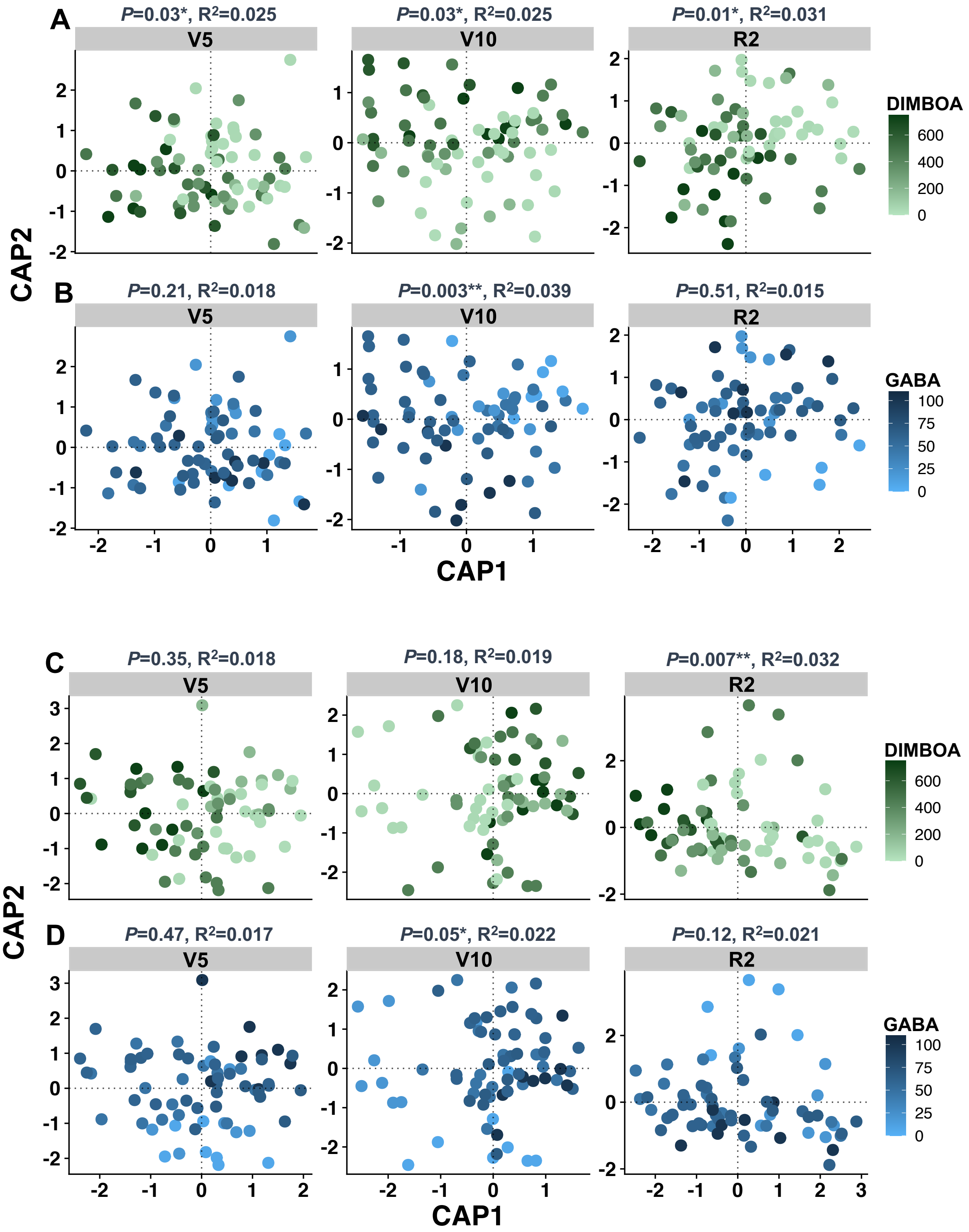


**Supplementary Figure S3. Effect of DIMBOA and GABA on the rhizosphere and root endosphere bacterial communities at different developmental stages including vegetative**

**stage 5 (V5), vegetative stage 10 (V10) and reproductive stage 2 (R2)**. (A) CAP analysis showing the DIMBOA effect on the bacterial communities in the rhizosphere at V5, V10, R2. (B) CAP showing the GABA effect on the bacterial communities in the rhizosphere at V5, V10, R2. (C) CAP analysis showing the DIMBOA effect on the bacterial communities in the root endosphere at V5, V10, R2. (D) CAP showing the GABA effect on the bacterial communities in the root endosphere at V5, V10, R2. PerMANOVA *p*-value and $R^2$ are displayed in each CAP plot and result from a model constraining DIMBOA and GABA, and factoring out the effect of experimental blocks. The quantitative data of DIMBOA and GABA concentration in root exudates collected from the semi-hydroponic system was used for the analysis. All the rhizosphere or root endosphere samples from the nine genotypes were separated into V5, V10 and R2 stages to perform the analysis.

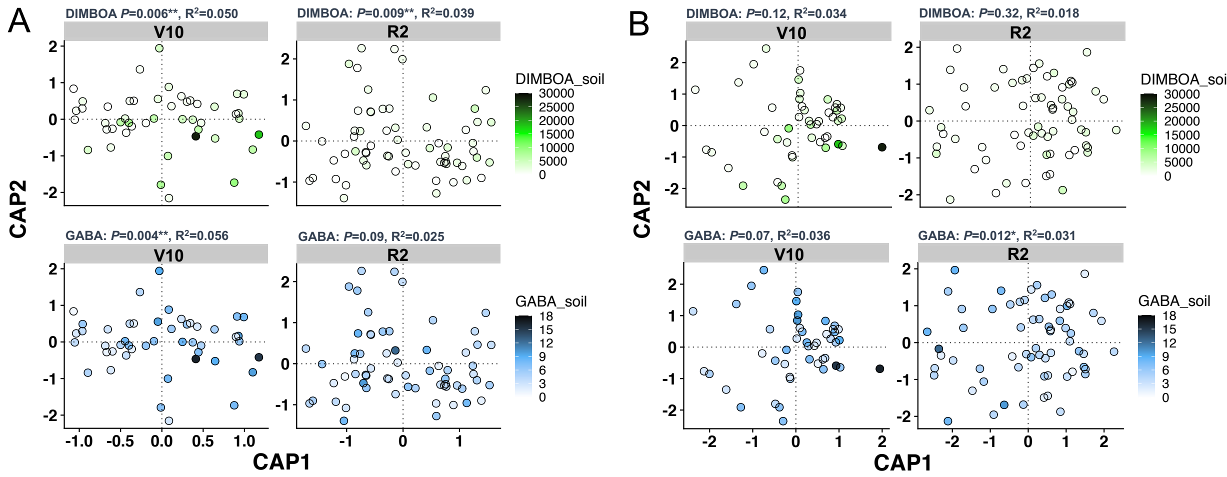


**Supplementary Figure S4. Quantification of DIMBOA and GABA in the rhizosphere soil collected from the field confirms the effects on the root-associated bacterial communities measured when using the exudate concentrations from the semi-hydroponic system.** CAP analysis showing that DIMBOA and GABA measured from the rhizosphere soil affects the rhizosphere and root endosphere bacterial communities in a similar pattern as using the data from the semi-hydroponic system. (A) CAP showing the effects of DIMBOA and GABA on the rhizosphere bacterial community structure at the V10 and R2 plant growth stages. (B) CAP showing the effects of DIMBOA and GABA on the root endosphere bacterial community structure at the V10 and R2 plant growth stages. PerMANOVA *p*-values and $R^2$ are displayed in each CAP plot and result from a model constraining DIMBOA and GABA, and factoring out the effect of experimental blocks. The quantitative data of DIMBOA and GABA concentration in the rhizosphere soil collected from the field was used for the analysis. All the rhizosphere or endosphere samples from nine genotypes were separated into V10 and R2 to perform the analysis.

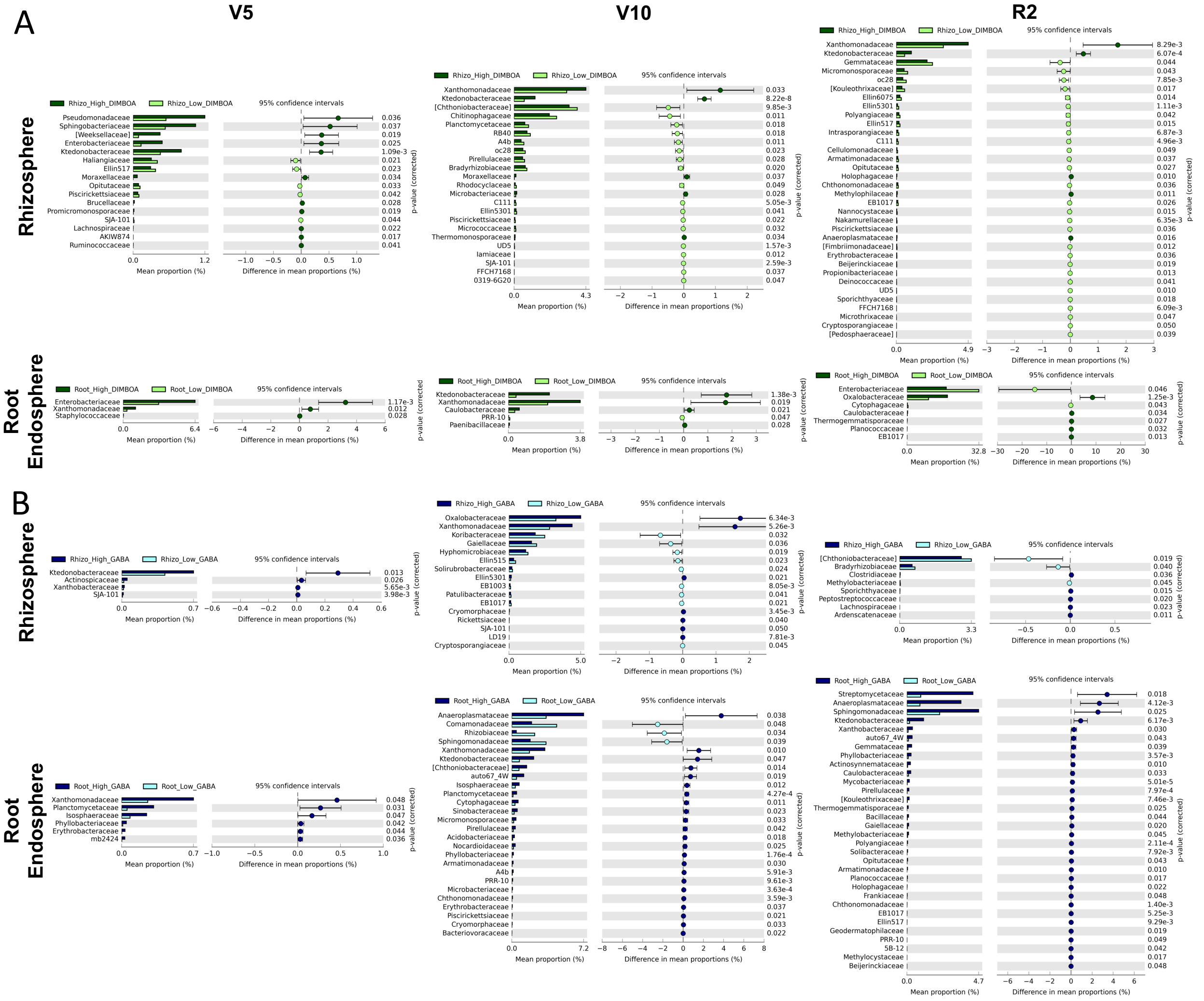


**Supplementary Figure S5. Changes in the relative abundance of specific bacterial families between maize genotypes exuding high and low DIMBOA or GABA levels at different plant growth stages.** (A) Bacterial families significantly affected by DIMBOA levels (high vs. low) in the root endosphere and rhizosphere at the V5, V10 and R2 developmental stages. (B) Bacterial families significantly affected by GABA levels (high vs. low) in the rhizosphere and root endosphere at the V5, V10 and R2 growth stages. The Welch's t-test was used to identify the bacterial families showing significant differences between DIMBOA and GABA levels on the software Statistical Analysis of Metagenomic Profiles (STAMP). Only the families with corrected

$p$-value $\leq 0.05$ (Bonferroni) were shown in the bar graphs. The classification of genotypes as low and high exuders of DIMBOA and GABA was according to Fig. 1.

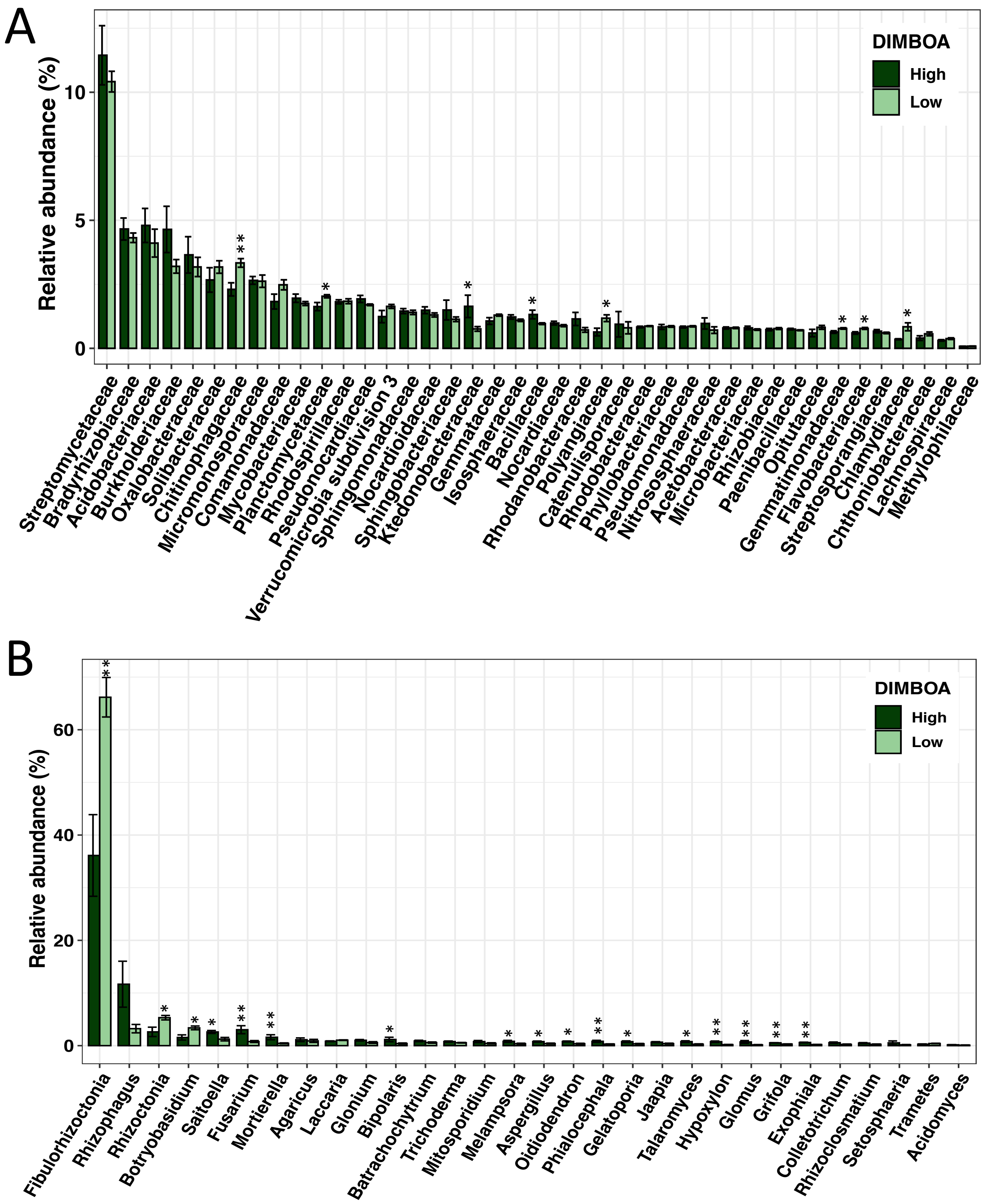


**Supplementary Figure S6. Shifts in the relative abundance of transcripts assigned to prokaryotic families and fungal genera between high and low DIMBOA levels in root**

**exudates at the V10 stage.** The bars reflect the proportion of transcript counts (gene expression levels) assigned to each specific microbial taxa based on the metatranscriptomics data. (A) Bacterial families changing in gene expression levels between genotypes exuding low and high DIMBOA in the root exudates. (B) Fungal genera changing in gene expression levels between genotypes exuding low and high DIMBOA in the root exudates. The top 41 most abundant bacterial families and 31 fungal genera were shown in the graphs. Only a subset of genotypes showing significant differences in the exudate concentration of DIMBOA were selected for the metatranscriptomics analysis. The maize lines PI587128, Ames26787 and Ames27171 were classified as low DIMBOA exuders, while PI550473 (B73) and Ames10248 were classified as high DIMBOA exuders. Each genotype has three replicates except B73 that has two replicates. Statistical analysis was done in the LEfSe software with Kruskal-Wallis test. The symbols "*" and "**" indicate significant differences at $p<0.05$ and $p<0.01$, respectively.
